# A chromosome-level genome of the franciscana dolphin, *Pontoporia blainvillei*

**DOI:** 10.64898/2026.07.31.742074

**Authors:** Lucas Eduardo Costa Canesin, Alexandre Aleixo, Amanda F. Vidal, Amely B. Martins, Ana Paula Cazerta Farro, Cristiane K. M. Kolesnikovas, Débora de Morais Cordeiro, Emanuel Bruno Neuhaus, Fábia O. Luna, Felipe A. A. Araújo, Gisele Nunes, Haydée A. Cunha, Izabela S. Mendes, Jacqueline S. de Mattos, Layse Albuquerque, Leandro Magalhães, Renato R. M. Oliveira, Sandro L. Bonatto, Silvia Britto Barreto, Daniel L. Z. Kantek, Sibelle T. Vilaça

## Abstract

The franciscana dolphin (Pontoporia *blainvillei*) is a small coastal cetacean endemic to the southwestern Atlantic Ocean and one of the most threatened marine mammals worldwide. It faces severe threats from bycatch, habitat degradation, and pollution. Classified as “Vulnerable” by the IUCN and “Critically Endangered” in Brazil, the species’ restricted range, strong fidelity to shallow waters, and low reproductive rate increase its extinction risk. Here, we present the first chromosome-level genome assembly for the franciscana dolphin, generated using PacBio HiFi long-read sequencing and Hi-C chromatin conformation capture. The final assembly totaled 3.13 Gb across 22 chromosomes (1500 scaffolds), consistent with the estimated karyotype of 2n = 44, with scaffold N50 of 111.18 Mb, high BUSCO completeness (99.42%), and a consensus quality value of 65.76. This high-quality genomic resource fills an important phylogenetic gap within Cetacea, enabling comparative and conservation studies. It provides an essential foundation for population genomics research to assess genetic diversity, structure, and connectivity, thereby supporting evidence-based conservation strategies for this endangered species.

## INTRODUCTION

*Pontoporia blainvillei* (Gervais and D’Orbigny, 1844), commonly known as franciscana dolphin, La Plata dolphin, or toninha in Brazil, is a small coastal odontocete endemic to the temperate and subtropical waters of the southwestern Atlantic Ocean. Its distribution ranges from the central Brazilian coast, between Itaúnas in Espírito Santo state southwards to the San Matías Gulf in Argentina (Crespo et al., 1998; Secchi et al., 2023) (Figure 1A). Recently the Society for Marine Mammalogy Committee on Taxonomy recognized its subdivision into two subspecies, *P. blainvillei blainvillei* in the south (Argentina to southern Rio de Janeiro, Brazil) and *P. blainvillei pukusi* in the northern part of the range (northern Rio de Janeiro to Espírito Santo) (Nara et al., 2024). As one of the most threatened cetacean species globally, *P. blainvillei* faces severe pressures from incidental bycatch by fishing gears, coastal habitat degradation, and marine pollution (Secchi et al., 2021). Its restricted distribution, strong fidelity to shallow coastal and estuarine habitats (≤30 m depth) (Crespo, 2018), and unfavorable life-history traits such as a low reproductive rate, an estimated generation time of 12 years (Taylor et al., 2007), and a maximum longevity of 21 years (Pinedo & Hohn, 2000), contribute to its high extinction risk. These combined factors have led the International Union for Conservation of Nature (IUCN) to classify the species as “Vulnerable”, while national Red List assessments classified it as critically endangered in Brazil, endangered in Argentina and vulnerable in Uruguay, underscoring the urgent need for conservation actions tailored to specific areas. The availability of high-quality genomic resources is thus of great relevance to inform management units and conservation priorities.

**Figure 1.**
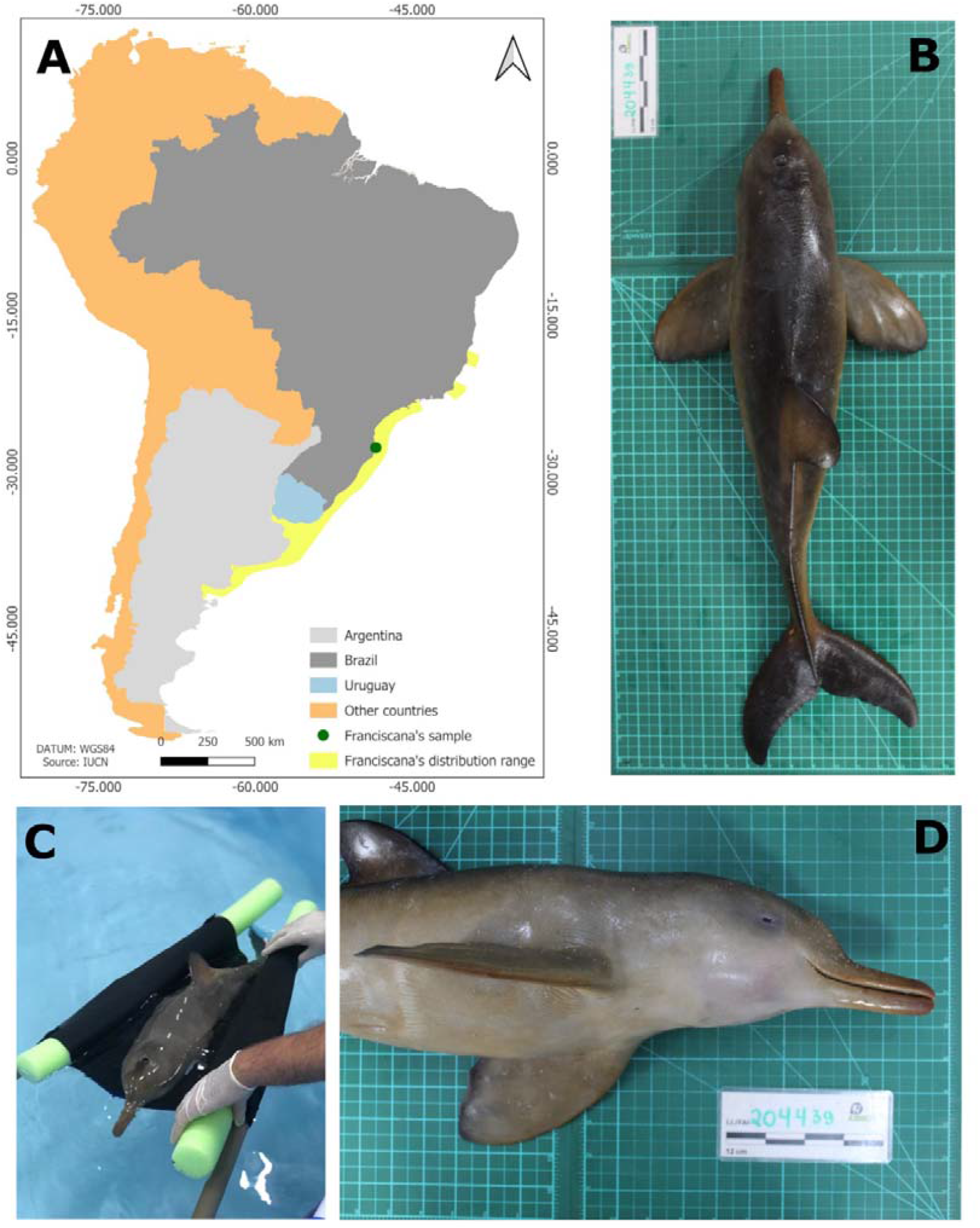
A) Distribution map of *Pontoporia blainvillei*. The green dot denotes the origin of the sampled and sequenced individual. C) Photo of sequenced specimen while in captivity. B) and D) Photo of the sequenced individual during sample collection (bar = 12cm).

Franciscana dolphins represent a unique evolutionary lineage, dating back to the Middle Miocene (approximately 20–15 million years ago). Molecular and fossil evidence strongly support its closest phylogenetic relationship with the Amazon River dolphin lineage (genus *Inia*) (Cassens et al., 2000; Hamilton, Caballero, Collins, & Brownell Jr, 2001). During the Miocene their ancestors inhabited epicontinental seas across South America, such as the Pebas System. As these seas regressed and modern river basins formed, the *Pontoporia* lineage became isolated within the Paraná Basin, following a distinct evolutionary trajectory. Eventually, it reached the Atlantic Ocean via the Río de la Plata estuary, initiating its coastal colonization (Cunha, 2022; Nara, Cremer, Farro, Colosio, Barbosa, Bertozzi, Secchi, Pagliani, Costa-Urrutia, & Gariboldi, 2022).

High-quality reference genomes are essential for comparative genomics, phylogenetics, and conservation biology, as they enable the identification of genes under selection, the detection of historical demographic events, and the assessment of gene flow among populations (Koepfli et al., 2015; Teixeira & Huber, 2021). Although draft genome assemblies have previously been generated for *P. blainvillei*, no chromosome-level assembly was available until now.

In this study, we present the first phased chromosome-level genome assembly of the franciscana dolphin, generated using PacBio HiFi long-read sequencing and Hi-C chromatin conformation capture. This high-quality reference genome provides a robust foundation for future studies on population structure, adaptive variation, and conservation-relevant genetic markers, facilitating the definition of evolutionary significant units and priority areas for protection. In the face of escalating threats to coastal ecosystems, this genomic resource represents a critical step toward improving the long-term survival prospects of this species.

## METHODS

### Sampling

Samples were collected on September 11, 2023, from a female of *P. blainvillei blainvillei* undergoing rehabilitation at the R3 Animal Rehabilitation Center in Florianópolis, Brazil (Figure 1B-D). The specimen, stranded at Praia da Joaquina, in the same city, died while under care at the facility. Biological material was collected postmortem and subsequently stored at –80 °C. Samples were transferred on dry ice to the Vale Institute of Technology (ITV), where they remained at –80 °C until further laboratory processing. Biological material was collected under permit 640/2015, issued by the Brazilian “Authorization for Capture, Collection, and Transport of Biological Material” (ABIO), and registered in the “National System for the Management of Genetic Heritage and Associated Traditional Knowledge” (SisGen) under code A88789E.

### Library preparation and sequencing

High-molecular weight DNA was extracted from a liver sample using the Nanobind Tissue Big DNA Kit (Pacific Biosciences). For long-read sequencing, a ~20 Kb library was prepared with the SMRTbell Express Template Prep Kit 2.0 and sequenced on a PacBio Sequel IIe using SMRT Cell 8M Tray.

A chromatin conformation capture (Hi-C) library was constructed to achieve chromosome-level assembly of the genome. Crosslinked and biotin-labeled DNA was obtained using the Arima Hi-C 2.0 Kit (Arima Genomics). The library was prepared with the NEBNext Ultra II DNA Library Prep Kit (New England Biolabs) from a kidney sample and sequenced on an Illumina NextSeq 2000 using a P3 Reagent 300-cycle kit (2 x 150 base pairs; bp).

RNA-seq libraries were generated for liver, kidney, heart, and central nervous system to support genome annotation. Total RNA was extracted using the RNeasy Mini Kit (Qiagen) and quality-assessed with Qubit (Thermo Fisher Scientific) and TapeStation (Agilent Technologies). RNA libraries were prepared using TruSeq Stranded Total RNA (Illumina) and sequenced on an Illumina NextSeq 2000 with a P3 Reagent 200-cycle kit (2 x 101 bp).

### Genome assembly and annotation

The genome was assembled using Pipeasm, an automated Snakemake pipeline that follows the best practices proposed by the Earth BioGenome Project (Marques Silva et al., 2026). A listing of tools used in the assembly process is outlined in Table 1. In summary, PacBio HiFi-CCS long reads and Hi-C short reads were trimmed using Cutadapt v4.4 (Martin, 2011) and fastp v0.23.4 (Chen et al., 2018), respectively, to ensure high-quality input data. Quality control was performed with FastQC v0.12.1 (Andrews, 2010) for both long and short reads, while long HiFi reads were also checked with NanoPlot v1.41.6 (De Coster & Rademakers, 2023). K-mer profiling was conducted with Meryl v1.3 (Rhie et al., 2020), and genome features (size, heterozygosity, and repeat content) were estimated using GenomeScope v2.0 (Ranallo-Benavidez et al., 2020). SmudgePlot v0.3.0 (Ranallo-Benavidez et al., 2020) was used to identify ploidy levels through K-mer distribution analysis, and KAT (Mapleson et al., 2017) was used to investigate possible GC content deviations in relation to K-mer distribution. The core assembly was generated with Hifiasm v0.19.6 (Cheng et al., 2021) using both HiFi and Hi-C reads. Assembly graphs were converted to FASTA and summarized with GFAstats v1.3.6 (Formenti et al., 2022). Post-assembly, the mitogenome was recovered with MitoHifi v3.2.2 (Uliano-Silva et al., 2023), and contaminants were removed using FCS Adaptor v0.5.0 and FCS-GX v0.5.0 (Astashyn et al., 2024). Genome completeness was assessed with Compleasm v0.2.2 using the mammalia_odb10 BUSCO dataset (Huang & Li, 2023; Manni et al., 2021). Merqury (Rhie et al., 2020) was used to evaluate K-mer-based assembly quality, and BlobToolkit v4.3.5 (Challis et al., 2020) generated snailplots summarizing assembly metrics.

**Table 1.**
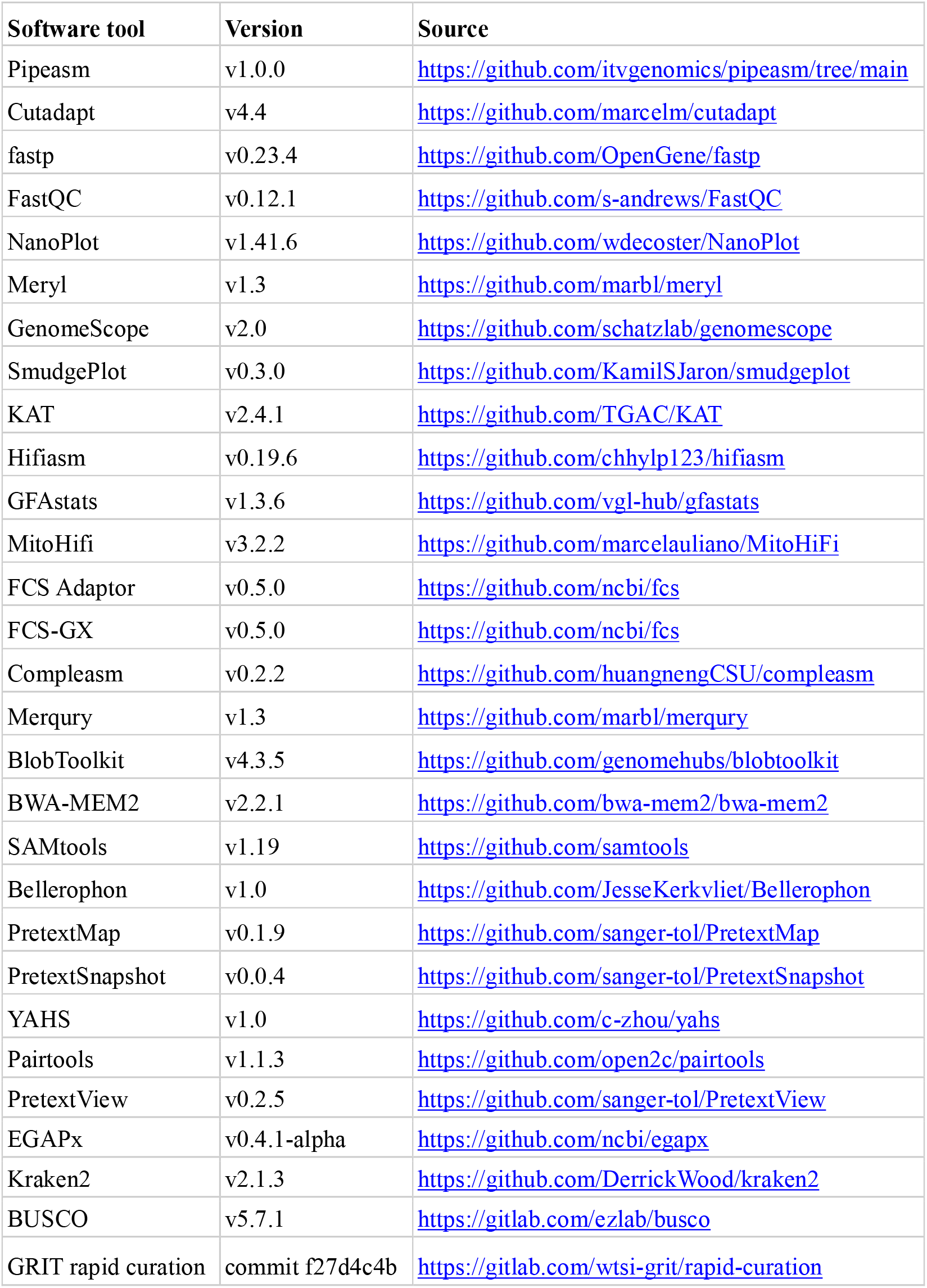
Software versions with respective sources, for all programs and pipelines that were used in this work.

To elucidate chromosomal interactions for scaffolding, reads from chromatin conformation capture libraries (Hi-C) were mapped to phased assemblies. Genome indexing and mapping for both haplotypes from decontaminated assemblies were performed using BWA-MEM2 v2.2.1 (Vasimuddin et al., 2019). Subsequently, forward and reverse Hi-C reads were aligned to each haplotype and sorted with SAMtools v1.19 (Li et al., 2009). Low-quality alignments and duplicated paired-end reads were filtered and merged using Bellerophon v1.0 (https://github.com/davebx/bellerophon). Quality assessment was performed using SAMtools flagstat and the script get_stats.pl (https://github.com/ArimaGenomics/mapping_pipeline). We used PretextMap v0.1.9 (https://github.com/sanger-tol/PretextMap) and PretextSnapshot v0.0.4 (https://github.com/sangertol/PretextSnapshot) for visual representations of Hi-C contact maps. Automatic scaffolding was carried out using YAHS v1.0 (Zhou et al., 2023), linking contigs into longer scaffolds to improve assembly contiguity. After automatic scaffolding, we manually curated the genome by remapping the Hi-C reads to each haplotype using BWA-MEM2 and filtered the resulting alignment using pairtools (Open2C et al., 2024), allowing the inclusion of repetitive sequence Hi-C signals. Manual curation of each haplotype was performed in PretextView 0.2.5 (https://github.com/sangertol/PretextView). Curated haplotypes were concatenated into a single assembly using GRIT Rapid Curation scripts (https://gitlab.com/wtsi-grit/rapid-curation). Curation steps were repeated on the concatenated assembly, followed by dual manual curation in PretextView. Finally, the assembly was split into two distinct haplotypes again using GRIT Rapid Curation scripts, and quality checks were performed on both curated phased haplotypes with GFAstats, Compleasm, and Merqury.

Gene prediction was performed on haplotype 1, which is the best assembly and includes the sexual chromosomes, using the NCBI Eukaryotic Genome Annotation Pipeline (https://github.com/ncbi/egapx). RNA-seq data from four tissues (liver, kidney, heart, and central nervous system) supported structural annotation. Reads were screened for bacterial contamination with Kraken2 (Wood et al., 2019), and any identified bacteria-derived reads were filtered out. Annotation completeness was evaluated with BUSCO (Simão et al., 2015).

## RESULTS

### Genome assembly and annotation

We generated a phased chromosome-level genome assembly for the franciscana dolphin, designated mPonBla1.0 (Figure 1; Table 2). We obtained 28.99 million PacBio CCS-HiFi reads, totaling 187.07 Gb (59.77x coverage), and 353 Gb of Hi-C paired reads (112.78x coverage). Initial genome size estimation, based on k-mer profile analysis of raw long-read data, indicated a size of 2.8 Gb and a heterozygosity of 0.2% (Figure 2A). No contamination was detected in the assembly (Figure 2B). The repetitive content was estimated as 29.3% of the total genome length. Both haplotypes were anchored into 22 chromosomes, consistent with its karyotype (2n = 44) (Figure 2C and 2D) (Heinzelmann et al., 2009). The curated reference haplotype (haplotype 1) spans 3.13 Gb (3,127,100,229 bp) and comprises 1,500 scaffolds (2,288 contigs), with a contig N50 of 17.36 Mb and scaffold N50 of 111.18 Mb (Figure 2E). The secondary assembly (haplotype 2) had a total size of 2.65 Gb (2,651,950,299 bp) across 1,215 scaffolds (1,824 contigs), with a contig N50 of 20.32 Mb and a scaffold N50 of 109.8 Mb (Figure 2F).

**Table 2.**
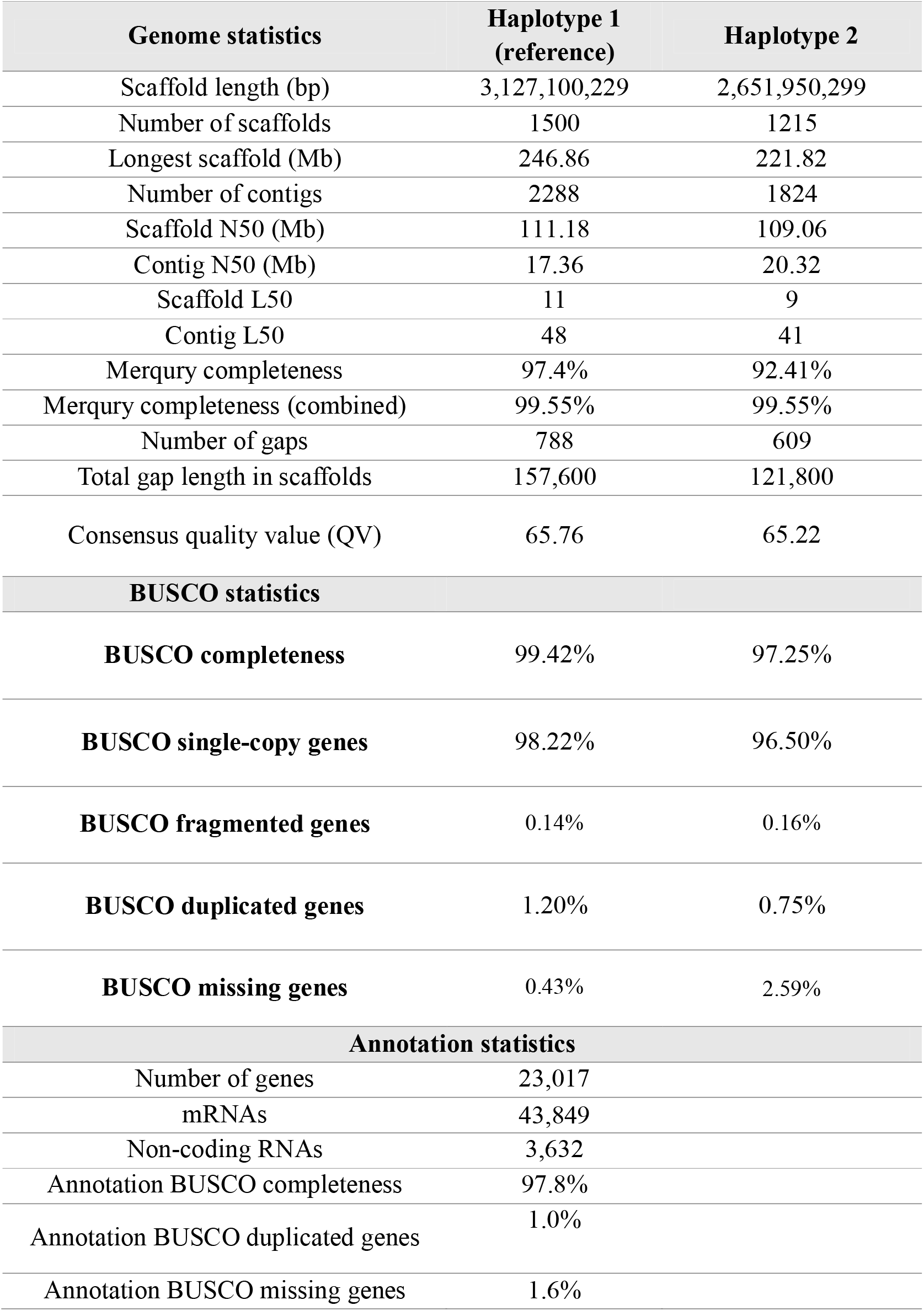
Genome assembly metrics for *Pontoporia blainvillei*.

**Figure 2.**
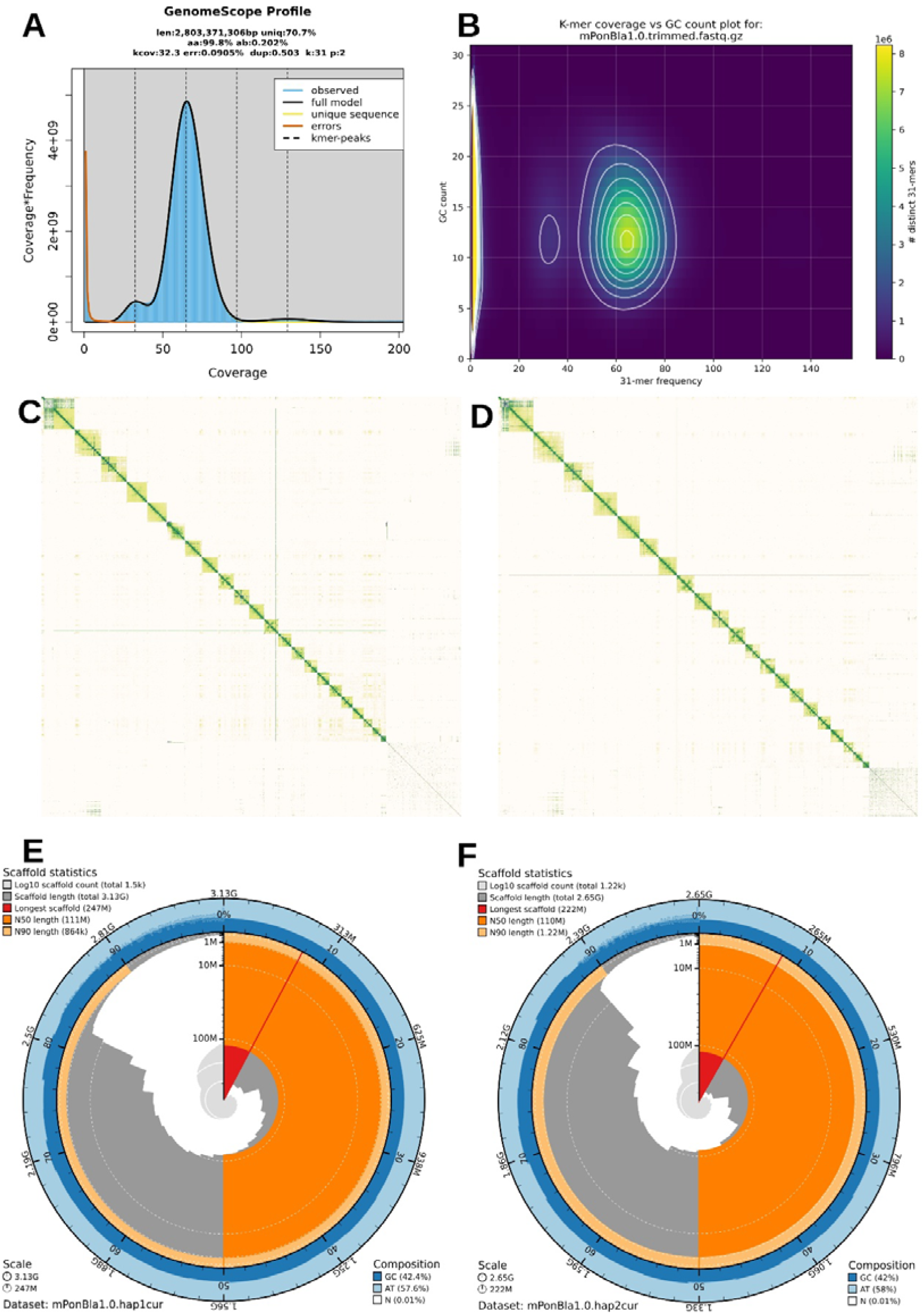
Genome assembly statistics and quality assessment of *Pontoporia blainvillei*. A) GenomeScope profile based on k-mer distribution. B) Density plot showing GC versus kmer content. C) HiC contact maps for primary and D) alternative assembly depicting the dual curation. E) Snailplot for haplotype 1 assembly showing an overview of assembly metrics and BUSCO gene completeness. F) Snailplot for haplotype 2 assembly.

Chromosome lengths ranged from 246.86 Mb to 49.7 Mb. The read k-mer completeness of haplotype 1 was 97.4%, with combined haplotypes reaching 99.55%, and the consensus quality value (QV) was 65.76 (Table 2). BUSCO analysis of the primary haplotype using the *mammalia_odb10* database revealed 99.42% completeness (98.22% single-copy, 1.20% duplicated). Haplotype 2 showed 96.5% completeness with 0.75% duplication. Compared to previously available *P. blainvillei* genomes, the assembly obtained in this study is the most complete and contiguous to date (Table 3).

**Table 3.** Comparison between statistics of previously available genomes for *Pontoporia blainvillei* and the one generated in this study.

|  | **This study**<br>**(haplotype 1)** | **GCA_011754075.1** | **GCA_004363935.1** |
| --- | --- | --- | --- |
| **Genome size (bp)** | 3,127,100,229 | 2,011,747,776 | 1,685,099,017 |
| **Total ungapped length**<br>**(bp)** | 3,126,942,629 | 2,006,438,987 | 1,685,034,717 |
| **Number of scaffolds** | 1,500 | 493,481 | 1,885,058 |
| **Scaffold N50 (bp)** | 111,176,753 | 493,481 | 2,541 |
| **Scaffold L50** | 11 | 85,936 | 134,511 |
| **Number of contigs** | 2,288 | 626,452 | 1,885,701 |
| **Contig N50 (bp)** | 17,358,612 | 5,888 | 2,541 |
| **Contig L50** | 48 | 100,720 | 135,152 |
| **GC percent (%)** | 42.39 | 41 | 46.5 |
| **Genome coverage** | 59.77x | 18x | 55.3x |
| **Sequencing technology** | PacBio HiFi +<br>Hi-C Illumina<br>NextSeq | Illumina HiSeq | Illumina HiSeq |

Gene prediction yielded 23,017 gene models, including 43,849 mRNAs and 3,632 non-coding RNAs (Table 2). Annotation quality was assessed using BUSCO with the *eutheria_odb10* database, showing 97.8% complete BUSCO genes, 1.0% duplicated, and 1.6% missing. These metrics demonstrate the high completeness and quality of the genome structural annotation (Table 2).

## DISCUSSION

Recent advances in genomics have established new standards for reference genome quality within the infraorder Cetacea (Morin et al., 2020). Chromosome-level assemblies for 18 cetacean species across eight families highlighted the importance of highly contiguous genomes for evolutionary and conservation research (Morin et al., 2025). The availability of a high-quality *P. blainvillei* genome fills a critical phylogenetic gap, as Pontoporiidae was previously unrepresented among contiguous assemblies.

The high contiguity and completeness of the *P. blainvillei* genome provide a robust foundation for downstream analyses. Our metrics meet the standards proposed by large-scale initiatives such as Earth Biogenome Project (Lawniczak et al., 2022) and were generated by the Genomics of the Brazilian Biodiversity (GBB) consortium, the largest national initiative dedicated to producing high-quality genomic data for Brazilian species, with a primary focus on supporting biodiversity conservation (Vilaça et al., 2024). Compared to previously available assemblies for this species, our chromosome-level genome represents a substantial improvement in contiguity and accuracy, enabling high-resolution studies of structural variation and adaptive evolution.

This resource also opens new opportunities for future studies in comparative genomics. *Pontoporia blainvillei* is the closest extant relative of the Amazon river dolphins of the genus *Inia*, for which a chromosome-level genome assembly (*I. geoffrensis*) is publicly available (Morin et al., 2025). These sister genera diverged during the Miocene and subsequently adapted to contrasting aquatic environments: *Inia* to freshwater river systems and *Pontoporia* to coastal marine habitats (Hamilton, Caballero, Collins, & Brownell, 2001; Nara, Cremer, Farro, Colosio, Barbosa, Bertozzi, Secchi, Pagliani, Costa-Urrutia, Gariboldi, et al., 2022). With chromosome-scale assemblies now available for both species, comparative genomic analyses can investigate chromosomal rearrangements, identify lineage-specific genes, detect signatures of positive selection, and explore the evolution of gene families and repetitive elements. Such studies will clarify molecular mechanisms underlying ecological divergence, refine the phylogenetic placement of both taxa within Odontoceti, and inform conservation strategies by providing insights into genome-wide patterns of variation and adaptation.

Finally, this chromosome-level genome will serve as a foundational resource for future population genomics studies, which are essential to generate data for assessing genetic diversity, structure, and connectivity among franciscana dolphin populations. Such information is critical for evidence-based conservation planning, particularly for a species facing severe anthropogenic threats and classified as Critically Endangered in Brazil (ICMBio, 2023). These efforts are aligned with the objectives of the Brazilian National Action Plan (PAN) for Toninha/Franciscana (https://www.gov.br/icmbio/pt-br/assuntos/biodiversidade/pan/pan-toninha), which aims to prevent population decline of the franciscana dolphin across all management areas, primarily through reducing incidental captures and protecting critical habitats. Genomic data will strengthen the implementation of this plan by supporting the identification of management units, knowledge of possible genetic erosion, and guiding long-term recovery actions.

## DATA AVAILABILITY

The chromosome-level genome assembly of *Pontoporia blainvillei* is publicly available through European Nucleotide Archive (ENA) accession numbers PRJEB122094 (ERP202127) for haplotype 1 and PRJEB122095 (ERP202128) for haplotype 2.

## ACKNOWLEDGEMENTS

This study was part of the Genomics of the Brazilian Biodiversity (GBB) consortium, a partnership between the Chico Mendes Institute for Biodiversity Conservation (ICMBio) and the Vale Institute of Technology (ITV), funded by Vale S.A. under the agreement “Acordo de Parceria PD&I Nº 01/2022”. Samples from the Associação R3 Animal were collected as part of the Santos Basin Beach Monitoring Project (PMP/BS), a requirement set by the Brazilian Institute of the Environment (IBAMA) for the environmental licensing of oil and natural gas production and transportation by Petrobras in the pre-salt province (25□05’S 42□35’W to 25□55’S 43□34’W) under ABIO Nº 640/2015. AA thanks the Brazilian Research Council (CNPq) for a research productivity fellowship (309243/2023-8).

## References

Andrews, S. (2010). FastQC: a quality control tool for high throughput sequence data. Babraham Bioinformatics, Babraham Institute, Cambridge, United Kingdom.

Astashyn, A., Tvedte, E. S., Sweeney, D., Sapojnikov, V., Bouk, N., Joukov, V., Mozes, E., Strope, P. K., Sylla, P. M., & Wagner, L. (2024). Rapid and sensitive detection of genome contamination at scale with FCS-GX. Genome Biology, 25(1), 60.

Cassens, I., Vicario, S., Waddell, V. G., Balchowsky, H., Van Belle, D., Ding, W., Fan, C., Mohan, R. S. L., Simoes-Lopes, P. C., & Bastida, R. (2000). Independent adaptation to riverine habitats allowed survival of ancient cetacean lineages. Proceedings of the National Academy of Sciences, 97(21), 11343–11347.

Challis, R., Richards, E., Rajan, J., Cochrane, G., & Blaxter, M. (2020). BlobToolKit– interactive quality assessment of genome assemblies. G3: Genes, Genomes, Genetics, 10(4), 1361–1374.

Cheng, H., Concepcion, G. T., Feng, X., Zhang, H., & Li, H. (2021). Haplotype-resolved de novo assembly using phased assembly graphs with hifiasm. Nature Methods, 18(2), 170–175.

Chen, S., Zhou, Y., Chen, Y., & Gu, J. (2018). fastp: an ultra-fast all-in-one FASTQ preprocessor. Bioinformatics, 34(17), i884–i890.

Crespo, E. A. (2018). Franciscana Dolphin. In Encyclopedia of Marine Mammals (pp. 388–392). Elsevier. 10.1016/B978-0-12-804327-1.00133-3

Crespo, E. A., Harris, G., & González, R. (1998). Group size and distributional range of the franciscana, Pontoporia blainvillei. Marine Mammal Science, 14(4), 845–849.

Cunha, H. A. (2022). Genetic diversity, population structure, and phylogeography. In The Franciscana Dolphin (pp. 111–126). Elsevier. 10.1016/B978-0-323-90974-7.00018-5

De Coster, W., & Rademakers, R. (2023). NanoPack2: population-scale evaluation of long-read sequencing data. Bioinformatics, 39(5), btad311.

Formenti, G., Abueg, L., Brajuka, A., Brajuka, N., Gallardo-Alba, C., Giani, A., Fedrigo, O., & Jarvis, E. D. (2022). Gfastats: conversion, evaluation and manipulation of genome sequences using assembly graphs. Bioinformatics, 38(17), 4214–4216.

Hamilton, H., Caballero, S., Collins, A. G., & Brownell Jr, R. L. (2001). Evolution of river dolphins. Proceedings of the Royal Society of London. Series B: Biological Sciences, 268(1466), 549–556.

Hamilton, H., Caballero, S., Collins, A. G., & Brownell, R. L. (2001). Evolution of river dolphins. Proceedings of the Royal Society of London. Series B: Biological Sciences, 268(1466), 549–556. 10.1098/rspb.2000.1385

Heinzelmann, L., Chagastelles, P. C., Danilewicz, D., Chies, J. A. B., & Andrades-Miranda, J. (2009). The karyotype of Franciscana dolphin (Pontoporia blainvillei). Journal of Heredity, 100(1), 119–122.

Huang, N., & Li, H. (2023). compleasm: a faster and more accurate reimplementation of BUSCO. Bioinformatics, 39(10), btad595.

ICMBio. (2023). Sistema de Avaliação do Risco de Extinção da Biodiversidade – SALVE. https://salve.icmbio.gov.br

Koepfli, K.-P., Paten, B., & O’Brien, S. J. (2015). The Genome 10K Project: A Way Forward. Annual Review of Animal Biosciences, 3(1), 57–111. 10.1146/annurev-animal-090414-014900

Lawniczak, M. K. N., Durbin, R., Flicek, P., Lindblad-Toh, K., Wei, X., Archibald, J. M., Baker, W. J., Belov, K., Blaxter, M. L., Marques Bonet, T., & others. (2022). Standards recommendations for the earth BioGenome project. Proceedings of the National Academy of Sciences, 119(4), e2115639118.

Li, H., Handsaker, B., Wysoker, A., Fennell, T., Ruan, J., Homer, N., Marth, G., Abecasis, G., & Durbin, R. (2009). The Sequence Alignment/Map format and SAMtools. Bioinformatics, 25(16), 2078–2079. 10.1093/bioinformatics/btp352

Manni, M., Berkeley, M. R., Seppey, M., Simão, F. A., & Zdobnov, E. M. (2021). BUSCO update: novel and streamlined workflows along with broader and deeper phylogenetic coverage for scoring of eukaryotic, prokaryotic, and viral genomes. Molecular Biology and Evolution, 38(10), 4647–4654.

Mapleson, D., Garcia Accinelli, G., Kettleborough, G., Wright, J., & Clavijo, B. J. (2017). KAT: a K-mer analysis toolkit to quality control NGS datasets and genome assemblies. Bioinformatics, 33(4), 574–576.

Marques Silva, B., Trindade, F. de J., Costa Canesin, L. E., Souza, G., Aleixo, A., Nunes, G., & Moreira-Oliveira, R. R. (2026). Pipeasm: a tool for automated large chromosome-scale genome assembly and evaluation. Bioinformatics Advances, 6(1), vbaf326.

Martin, M. (2011). Cutadapt removes adapter sequences from high-throughput sequencing reads. EMBnet.Journal, 17(1), 10–12. 10.14806/ej.17.1.200

Morin, P. A., Alexander, A., Blaxter, M., Caballero, S., Fedrigo, O., Fontaine, M. C., Foote, A. D., Kuraku, S., Maloney, B., McCarthy, M. L., McGowen, M. R., Mountcastle, J., Nery, M. F., Olsen, M. T., Rosel, P. E., & Jarvis, E. D. (2020). Building genomic infrastructure: Sequencing platinumlJstandard referencelJquality genomes of all cetacean species. Marine Mammal Science, 36(4), 1356–1366. 10.1111/mms.12721

Morin, P. A., Bein, B., Bortoluzzi, C., Bukhman, Y. V., Hains, T., Heimeier, D., Uliano-Silva, M., Absolon, D. E., Abueg, L., Antosiewicz-Bourget, J., Balacco, J. R., Bonde, R. K., Brajuka, N., Brownlow, A. C., Carroll, E. L., Carter, M., Collins, J., Davison, N. J., Denton, A., … Jarvis, E. D. (2025). Genomic infrastructure for cetacean research and conservation: reference genomes for eight families spanning the cetacean tree of life. Frontiers in Marine Science, 12. 10.3389/fmars.2025.1562045

Nara, L., Cremer, M. J., Farro, A. P. C., Colosio, A. C., Barbosa, L. A., Bertozzi, C. P., Secchi, E. R., Pagliani, B., Costa-Urrutia, P., & Gariboldi, M. C. (2022). Phylogeography of the endangered franciscana dolphin: timing and geological setting of the evolution of populations. Journal of Mammalian Evolution, 29(3), 609–625.

Nara, L., Cremer, M. J., Farro, A. P. C., Colosio, A. C., Barbosa, L. A., Bertozzi, C. P., Secchi, E. R., Pagliani, B., Costa-Urrutia, P., Gariboldi, M. C., Lazoski, C., & Cunha, H. A. (2022). Phylogeography of the Endangered Franciscana Dolphin: Timing and Geological Setting of the Evolution of Populations. Journal of Mammalian Evolution, 29(3), 609–625. 10.1007/s10914-022-09607-7

Nara, L., Secchi, E. R., & Cunha, H. A. (2024). Divergence, diagnosability, and description of a new subspecies of franciscana dolphin Pontoporia blainvillei (Gervais & d’Orbigny, 1844). Journal of Mammalian Evolution, 31(3), 32.

Open2C, Abdennur, N., Fudenberg, G., Flyamer, I. M., Galitsyna, A. A., Goloborodko, A., Imakaev, M., & Venev, S. V. (2024). Pairtools: from sequencing data to chromosome contacts. PLOS Computational Biology, 20(5), e1012164.

Pinedo, M. C., & Hohn, A. A. (2000). Growth layer patterns in teeth from the franciscana, Pontoporia blainvillei: developing a model for precision in age estimation. Marine Mammal Science, 16(1), 1–27.

Ranallo-Benavidez, T. R., Jaron, K. S., & Schatz, M. C. (2020). GenomeScope 2.0 and Smudgeplot for reference-free profiling of polyploid genomes. Nature Communications, 11(1), 1432.

Rhie, A., Walenz, B. P., Koren, S., & Phillippy, A. M. (2020). Merqury: reference-free quality, completeness, and phasing assessment for genome assemblies. Genome Biology, 21(1), 1–27.

Secchi, E. R., Cremer, M. J., Danilewicz, D., & Lailson-Brito, J. (2021). A Synthesis of the Ecology, Human-Related Threats and Conservation Perspectives for the Endangered Franciscana Dolphin. Frontiers in Marine Science, 8. 10.3389/fmars.2021.617956

Secchi, E. R., Danilewicz, D., & Ott, P. H. (2023). Applying the phylogeographic concept to identify franciscana dolphin stocks: implications to meet management objectives. J. Cetacean Res. Manage., 5(1), 61–68. 10.47536/jcrm.v5i1.827

Simão, F. A., Waterhouse, R. M., Ioannidis, P., Kriventseva, E. V., & Zdobnov, E. M. (2015). BUSCO: Assessing genome assembly and annotation completeness with single-copy orthologs. Bioinformatics, 31(19), 3210–3212. 10.1093/bioinformatics/btv351

Taylor, B., Chivers, S., Larese, J., & Perrin, W. (2007). Generation length and percent mature estimates for IUCN assessments of cetaceans. NOAA, NMFS, Southwest Fisheries Science Center Administrative Report LJ-07-01.

Teixeira, J. C., & Huber, C. D. (2021). The inflated significance of neutral genetic diversity in conservation genetics. Proceedings of the National Academy of Sciences, 118(10). 10.1073/pnas.2015096118

Uliano-Silva, M., Ferreira, J. G., Krasheninnikova, K., Formenti, G., Abueg, L., Torrance, J., Myers, E. W., Durbin, R., Blaxter, M., & McCarthy, S. A. (2023). MitoHiFi: a python pipeline for mitochondrial genome assembly from PacBio High Fidelity reads. BMC Bioinformatics, 24(1), 1–13.

Vasimuddin, M., Misra, S., Li, H., & Aluru, S. (2019). Efficient architecture-aware acceleration of BWA-MEM for multicore systems. 2019 IEEE International Parallel and Distributed Processing Symposium (IPDPS), 314–324.

Vilaça, S. T., Vidal, A. F., Pavan, A. C. D., Silva, B. M., Carvalho, C. S., Povill, C., Luna-Lucena, D., Nunes, G. L., Figueiró, H. V., Mendes, I. S., Bittencourt, J. A. P., Côrtes, L. G., Costa Canesin, L. E., Oliveira, R. R. M., Damasceno, R. P., Vasconcelos, S., Barreto, S. B., Tavares, V., Oliveira, G., … Aleixo, A. (2024). Leveraging genomes to support conservation and bioeconomy policies in a megadiverse country. Cell Genomics. 10.1016/j.xgen.2024.100678

Wood, D. E., Lu, J., & Langmead, B. (2019). Improved metagenomic analysis with Kraken 2. Genome Biology, 20(1), 257.

Zhou, C., McCarthy, S. A., & Durbin, R. (2023). YaHS: yet another Hi-C scaffolding tool. Bioinformatics, 39(1), btac808.

